# Phylogenetic Conflict in Bears Identified by Automated Discovery of Transposable Element Insertions in Low Coverage Genomes

**DOI:** 10.1101/123901

**Authors:** Fritjof Lammers, Susanne Gallus, Axel Janke, Maria A Nilsson

**Affiliations:** Senckenberg Biodiversity and Climate Research Centre, Senckenberg Gesellschaft für Naturforschung, Senckenberganlage 25, 60325 Frankfurt am Main, Germany.; Goethe University Frankfurt, Institute for Ecology, Evolution & Diversity, Biologicum, Max-von-Laue-Str.13, 60439 Frankfurt am Main, Germany.

**Author notes:** Corresponding author: Dr. Maria A. Nilsson Senckenberg Biodiversity and Climate Research Centre, Senckenberg Gesellschaft für Naturforschung, Senckenberganlage 25, 60325 Frankfurt am Main, Germany.

**Keywords:** Retrotransposition, bears, Ursidae, phylogeny, evolution, transposable elements

## Abstract

Compared to sequence analyses, phylogenetic reconstruction from transposable elements (TEs) offers an additional perspective to study evolutionary processes. However, detecting phylogenetically informative TE insertions requires tedious experimental work, limiting the power of phylogenetic inference. Here, we analyzed the genomes of seven bear species using high throughput sequencing data to detect thousands of TE insertions. The newly developed pipeline for TE detection called TeddyPi (TE detection and discovery for Phylogenetic Inference) obtained 150,513 high-quality TE insertions in the genomes of ursine and tremarctine bears. By integrating different TE insertion callers and using a stringent filtering approach, the TeddyPi pipeline produced highly reliable TE insertion calls, which were confirmed by extensive *in vitro* validation experiments. Screening for single nucleotide substitutions in the flanking regions of the TEs show that these substitutions correlate with the phylogenetic signal from the TE insertions. Our phylogenomic analyses show that TEs are a major driver of genomic variation in bears and enabled phylogenetic reconstruction of a well-resolved species tree, even with strong signals for incomplete lineage sorting and introgression. The analyses show that the Asiatic black, sun and sloth bear form a monophyletic clade. TeddyPi is open source and can be adapted to various TE and structural variation callers. The pipeline makes it easy to confidently extract thousands of TE insertions even from low coverage genomes of non-model organisms, opening new possibilities for biologists to study phylogenies, evolutionary processes as well as rates and patterns of (retro-)transposition and structural variation.

## Introduction

In a innovative analysis almost 20 years ago, rare genomic changes were used to confirm the close relationship between hippopotamus (Artiodactyla) and whales (Cetacea) (Shimamura et al. 1997; Nikaido et al. 1999). Transposable element (TE) insertions are a type of rare genomic changes that propagates in the genome via copy-and-paste (retrotransposons) or cut-and-paste (DNA transposons) mechanisms. Germline transposition events will be passed on to the descendants, making it possible to deduce phylogenetic relationships (Shimamura et al. 1997; Nikaido et al. 1999). In contrast to nucleotide substitutions which are prone to homoplasy by parallelisms, convergence and reversals, TE insertions are virtually homoplasy free. Parallel integration of TE insertions in the same loci in different species is highly improbable due to low germline insertion rates and the presence of different active TE families (Ray and Xing 2006). Also, the exact removal of TE insertions is very rare and usually leaves a detectable genetic ‘scar’ (van de Lagemaat et al. 2005). These features were very valuable for understanding of deep or complex divergences, like the early radiation of mammals and birds (Churakov et al. 2009; Nishihara et al. 2009; Hallström and Janke 2010; Suh et al. 2015).

Detecting phylogenetically informative TE insertions was initially challenging, because fully sequenced genomes were not available (Shimamura et al. 1997; Nikaido et al. 1999). Therefore, only experimental work could identify candidate TE loci of which only a minor fraction was phylogenetically informative (Shimamura et al. 1997; Nikaido et al. 1999). With increasing availability of genome assemblies, new methods allowed computational identification of phylogenetically informative TE insertions but extending the taxon sampling for species without available genomes relied still on experimental work (Kriegs et al. 2006; Churakov et al. 2009). Other methods that are not based on genome assemblies were limited in number of informative TE insertions that can be identified (Suh et al. 2012; Kuramoto et al. 2015). Finally, experimental enrichment protocols for TE insertions can identify thousands of informative loci, but require knowledge of the TE sequence and are biased towards loci with present TE insertions (Platt et al. 2015). A recently developed bioinformatic approach to detect novel TE insertions is to use the information from discordantly mapped paired-end short reads and does not require a *de novo* genome assembly for each species (Medvedev et al. 2009). Such ‘TE calling’ methods allowed studying TE insertion dynamics and other structural variations (SV) on a population-scale (Hormozdiari et al. 2013; Sudmant et al. 2015). This approach has also been applied to the great apes and to mice (Nellåker et al. 2012; Hormozdiari et al. 2013) showing its potential for phylogenetic inference. However, no phylogenetic study has applied TE calling methods to non-model organisms yet, for which often only draft genome assemblies and low coverage re-sequencing data are available.

In phylogenetics, a long-standing question is the evolutionary history of bears (Ursidae) that is differently reconstructed when mitochondrial, autosomal and gonosomal DNA sequences were analyzed, revealing high levels of phylogenetic discordance (Krause et al. 2008; Pagès et al. 2008; Hailer et al. 2012; Miller et al. 2012; Bidon et al. 2014; Kutschera et al. 2014). This phylogenetic incongruence among bears can be caused by introgressive hybridization and incomplete lineage sorting (ILS) (Maddison 1997), making analysis of genome-wide data necessary to understand these complex processes (Delsuc et al. 2005). However, the lack of whole genome sequences inhibited efficient screening for phylogenetically informative TE insertion events until the polar bear (*Ursus maritimus*) genome sequence and genome data of all other bear species became available (Miller et al. 2012; Liu et al. 2014; Kumar et al. 2016). These new genome data allows to detect TE insertions as additional independent phylogenomic markers to study the evolution of Ursidae. We developed the TeddyPi (TE detection and discovery for phylogenetic inference) pipeline to process data from TE and SV callers. TeddyPi pursues the idea of integrating different TE callers (Lin et al. 2015; Nelson et al. 2016) and extends it to routinely integrate TE insertion datasets from multiple samples to track integrations of TEs in orthologous loci and to create presence/absence tables for phylogenetic inference. The general effectiveness of TeddyPi and its reliability for extracting TE insertions from low-coverage genomes in non-model organisms was evaluated using all bears of the Ursine subfamily and the monotypic Tremarctinae. Studying the evolution of bears has the advantage that every species is represented by at least one genome, that their genomes are TE rich (about 40 %) and that bears evolved less than 5 million years ago (Ma). This allows to also observe nucleotide substitutions in the flanks around the TE insertions, that were mutationally saturated in deeper divergences. In addition, a nucleotide-based genome-wide phylogeny (Kumar et al. 2016) allows to compare nucleotide and TE-based phylogenomic reconstructions. The TeddyPi pipeline extracted an extensive catalog of 150,513 TE insertions to reconstruct the first TE-derived species tree of bears and which reveals varying rates of TE accumulation in their genomes.

## Materials and Methods

### Taxon sampling and genome sequencing

Illumina HiSeq generated whole genome sequencing data from Kumar et al. (2016) for six ursine bear species and the spectacled bear (*Tremarctos ornatus*) were obtained. For mapping, reads were quality-trimmed with Trimmomatic (Bolger et al. 2014) mapped with BWA (Li and Durbin 2010), duplicates reads were marked. In total, nine genomes with a mean coverage of 13.7X from seven species were analyzed (**Supplementary Table 1**). In comparison to the giant panda genome sequence (ailMel1), the polar bear genome sequence (Liu et al. 2014) has higher contiguity and contains potentially better assembled repeats because it is based on longer reads and was therefore the preferred choice for reference mapping.

### Considerations for nested reference genomes

Programs to detect TE insertions (in analogy to SNP callers named TE callers) depend on pairwise comparison between the paired-end short reads of a sample and the reference genome the reads were mapped to. Because most published TE callers can only detect nonreference (Ref-) TE insertions it is beneficial to have a reference genome that is phylogentically placed as outgroup to the taxa under study to be able to detect insertions across the complete phylogeny (**Supplementary Fig. 1**). If this is not possible, the use of only non-reference TE callers will lead to unresolved internodes and a skewed phylogenetic interpretation. For example, when the reference genome is nested inside the ingroup/tree, only TE insertions on the terminal branches are detectable or certain internodes cannot be resolved (**Supplementary Fig. 1**). To overcome such a bias, reference (Ref+) TE insertions, i.e. those shared with the reference genome need to be considered.

### Analysis of TEs in the polar bear genome sequence

Repetitive elements in the polar bear genome were identified using RepeatMasker in strict mode searching for carnivore-specific repeats (Repbase version 20140131). The script createRepeatLandscape.pl provided with RepeatMasker was used to calculate the repeat landscape. The LINE1 ORF2 sequence was retrieved from a full-length LINE1 found on the polar bear Y chromosome (Bidon et al. 2015) and used as BLAST query against the polar bear genome sequence (Altschul et al. 1990). Hits were filtered for full length, coding ORF2 copies and a maximum of three mismatches. Then, these sequences plus 7,000 bp flanking sequence on 5’ and 3’ ends were extracted from the polar bear genome sequence. Within these sequences a BLAST search for a coding LINE1 ORF1 sequence was performed to find LINE1 copies containing two coding ORFs. As additional proxy for LINE1 activity, we screened the polar bear and giant panda genome for the U6 snRNA (Accession No: M14486.1) using BLAST. According to Doucet et al. (2015) all hits with more than 97.5% identity, 26 bp alignment length and an E-value of < 10 were considered as full length hits. Additionally, we annotated 146,268 gaps totaling to 38 mega base pairs (Mb) in the polar bear genome; the majority of these gaps (138,041) were larger than 1 base pairs (bp).

### Detection of non-reference (Ref-) TE insertions

Reference mapped short reads were processed with RetroSeq (Keane et al. 2013) and Mobster (Thung et al. 2014) to identify insertions that are present in the corresponding genome while being absent in the reference genome. For RetroSeq, a minimum mapping quality of 30 was used, a TE mapping identity of 90% at 50% length. The upper coverage threshold was set to 2.5X of the samples’ sequencing depth. Mobster was run with default settings. A consensus library of 593 carnivore specific TEs was supplied to both programs to identify reads that match TE sequence and thus give information on the type of TE that has integrated. In addition, RetroSeq identifies reads matching the RepeatMasker track in the reference genome. Using the TeddyPi pipeline, callsets from RetroSeq and Mobster were filtered for calls falling in regions of undetermined bases (N) in the polar bear genome plus a window of 200 bp. Calls with less than 5 supporting reads were filtered as were any calls within 100 bp from annotated TEs of the same type in the polar bear genome. For stringency, both datasets were also masked for regions that had a depth of coverage below ⅓ or 2.5 times the mean coverage of the respective sample. Only overlapping calls from both programs were further utilized.

### Detection of reference insertion (Ref+) TE insertions

To detect TE insertions absent from at least one of the low coverage bear genomes and present in polar bear reference genome, Pindel (Ye et al. 2009) and Breakdancer (Chen et al. 2009) were utilized to mine the genomes for deletions, that are indicative for insertions in the reference genome (Nellåker et al. 2012). Pindel uses split-read (SR) information to obtain breakpoint information at a single-nucleotide level resolution. Because BreakDancer does not utilize SRs for SV-calling, start- and end-coordinates from deletions were used. BreakDancer was called using a maximum variant size of 10 kb and requiring at least five supporting reads to make a SV call. Pindel was run with the following parameters ‐‐report_interchromosomal_events false, ‐‐anchor_quality 30, -w 40. Only deletions were considered for further processing. For each sample, book-ended calls and overlapping calls were merged, filtered for N-regions in the reference genome within 200 bp flanking each call and for calls falling within tandem repeats in the reference (+ 50 bp flanking sequence). All calls in regions with a depth of coverage below 0.33X or 2.5X higher than average were excluded. The calls from Pindel and Breakdancer were merged to a non-redundant set. The start/end coordinates or the breakpoint of the deletion plus a window of +-50 bp, depending on which information was available, were used to detect intersection with annotated repeats in the polar bear reference genome. Deletion calls that matched duplicate RepeatMasker hits and appeared twice, were merged. When coordinates overlapped with more than one TE in the reference genome including a recent SINE insertion (i.e. SINEC_Ame subfamily) and the other TEs were not elements known to be active within Carnivora, it was called as SINE derived. If coordinates overlapped with different types of annotated TEs, and more than one was potentially active the event was recorded as ‘complex’. Predicted deletion loci between samples were attributed to the same locus if both were intersecting with a reference TE and the distance between their breakpoints was less than to 100 bp. To obtain reference insertion (Ref+) calls, presence/absence information was inverted, because our processed deletion calls reflect TE insertions occurred in the lineage leading to the reference genome. / (**Supplementary Fig 2**).

### Integration of Ref+ and Ref+ call sets, filtering and processing

To combine insertion and deletion datasets, results were integrated across all species. This module of TeddyPi (tpi_ortho.py) loads the final call sets for all species, internally sorts these by position, and merges overlapping and book-ended calls if not done before. Then BedTools window is called via pybedtools to create a presence/absence matrix (coded as 1 and 0 respectively) over all variants and taxa (variant × taxa) (Quinlan and Hall 2010; Dale et al. 2011).

Because breakpoint estimates might differ slightly between different taxa although originating from the same insertion event a final merging step was performed by Bedtools merge. Overlapping, book-ended, and events being apart up to 100 bp were merged. Presence/absence information from deletion calls was inverted (1 ↔ 0) to obtain reference insertions (Ref+) calls. The state of TE insertions in the reference genome was added with either 1 or 0 for Ref+ and Ref– events, respectively. Call sets for Ref+ and Ref– were saved as tab-separated file and converted to a NEXUS character matrix using the python-nexus package (Greenhill S. unpublished).

### Merging Ref+ and Ref− callsets, correcting for missing data

Ref+ and Ref− datasets were merged in the tpi_unite.py module of TeddyPi, and a final presence/absence matrix was created. A synthetic outgroup with state ‘0’ for all loci was added. For the Ref− dataset, loci that did not meet coverage criteria in all samples, were coded as missing data (“?” in the NEXUS matrix) for the sample with insufficient or excessive depth of coverage. The criteria were set for each sample individually to include only loci with coverage between 0.33X and 2.5X of the samples mean coverage.

### Phylogenetic inference from TE insertion calls

We processed SINE and LINE1 callsets separately and created Dollo parsimony trees in PAUP* (Swofford 2002)using the heuristic search with 500 replicates. Bootstrap support was calculated from 1,000 replicates. The trees were rooted using the synthetic outgroup. The number of SINE insertions for species-tree congruent and alternative topologies wereobtained from the presence/absence matrices and analyzed using the KKSC-test that conceptually transfers the D-statistics to TE insertion data (Durand et al. 2011; Kuritzin et al. 2016). Median networks for SINE insertions were calculated in SplitsTree 4 (Huson and Bryant 2006). Also, phylogenetic networks for Ref+ and Ref− data were calculated separately using all SINEs and LINE1s.

### Estimating TE insertion rates

SINE and LINE insertion counts were extracted from the parsimony-tree branch lengths and were divided by the branch times (in million years, Myr) estimated previously (Kumar et al. 2016) to get estimates on the relative insertion rate. To estimate per-generation insertion rates, the generation time for polar and brown bear was assumed to be 10 years (Tallmon et al. 2004; Cronin et al. 2009) and 6 years for the other bear species (Onorato et al. 2004; Kutschera et al. 2014).

### Genomic context of TE insertions

The genomic context the TE insertions was evaluated using the genome annotation from the polar bear genome (Liu et al. 2014). The TE insertion catalogue was screened for overlaps with 3’ and 5’ UTRs, introns, exons and intergenic regions.

### Flanking sequence analysis of TE insertion loci

To investigate the sequence variation around TE insertion sites, consensus sequence alignments were created using substitution calls from Kumar et al. (2016). First, 10 kb sequence up- and downstream of the insertion site were extracted and the maximum likelihood (ML) phylogeny was inferred with RaxML (Stamatakis 2014) for each flank and the concatenated sequence of both flanks. For automation and calling RAxML, the Dendropy package was utilized (Sukumaran and Holder 2010). To account for the possibly misaligned reads around the insertion site, the first 500 bp on each side of the insertion site were excluded. The question was, whether the flanking sequence yields the same phylogenetic signal as the presence/absence pattern of the TE insertion. Therefore, we checked if the species carrying the TE insertion form a monophyletic group in the ML-trees using the ETE toolkit (Huerta-Cepas et al. 2016). Furthermore, to gain insight in the phylogenetic signal in the TE flanking region a sliding window approach was applied to the same 10 kb flanking regions using non-overlapping 1 kb windows. For each window, sites were counted showing the same phylogenetic signal like the TE insertion pattern and divided by the number of segregating sites.

### Experimental validation screening

From the *in silico* dataset, loci were randomly selected for experimental verification. DNA samples from all ursine bears and the spectacled bear were included. For the Asian bear species and the spectacled bear the same DNA samples as for the Illumina genome sequencing were used for validation. We selected loci containing TE insertions supporting different topologies (**Supplementary Table 2**), including topologies in conflict with the species tree (e.g. presence in American black and Asiatic black bear or American black bear and sun bear).

For primer design, consensus sequence alignments from Kumar et al. (2016) were extracted, that spanned 4 kb up- and downstream of the predicted TE insertion site. PCR primers were generated with primer3 to be located approximately 200 bp from the TE insertion site(Untergasser et al. 2012). Primers are listed in the supplement (**Supplementary Data 1**). Each locus was amplified using 8 ng of DNA per species and Amplicon Taq (VWR) in a touchdown PCR. Banding patterns were examined using gel-electrophoresis agarose gels along with a 1 kb DNA marker (ThermoFisher GeneRuler 1Kb). The fragment length of each PCR product was estimated and species that had the indication of a TE insertion were recorded. The PCR amplicons were Sanger-sequenced in both directions using the ABI 3730 DNA Analyzer. Using the Sanger-sequenced TE-locus the type of the integration was determined by querying the sequence against Repbase (Jurka et al. 2005) (www.girinst.org). For 13 markers we sequenced the complete or near complete taxon-sampling to verify the phylogenetic information of the loci. The alignments were screened for the type of the TE, the orientation of the TE, TSDs and the integrity of the flank. One marker was specifically selected and sequenced to investigate the absence of a SINEC1_Ame in the polar bear (marker 40). The sequence analysis showed that the SINEC1_Ame, was missing in the polar bear.

Experimentally confirmed insertion patterns were compared with the computationally predicted insertions at the same locus. We considered each matching insertion status (predicted: absence - PCR: absence/ predicted: presence - PCR: presence) as correctly called. If the PCR product indicated presence of a TE insertion but no TE call was made, the locus was recorded as false negative (FN) and false positive (FP) for the opposite case. If a PCR reaction did not yield an amplicon for a locus, the locus was flagged as inconclusive.

## Results

### Transposable elements in ursine bears

Our screening of the interspersed repeats in the polar bear reference genome identified 1,223,168 SINEs (8.4%), 978,888 LINEs (21.3%), 320,346 LTR retrotransposons (5.3%), as well as 340,447 DNA transposons (3.1%) (**Supplementary Table 3**). In total, the polar bear genome is comprised by 38.1% interspersed repeats, similar to other carnivores like panda, dog or cat (Lindblad-Toh et al. 2005; Pontius et al. 2007; Li et al. 2010). The most abundant and recently active SINE-family in carnivore genomes is the Lysine-tRNA derived SINEC (Walters-Conte et al. 2011). In Ursidae, SINEC1_AMe is the most frequent SINE subfamily in both the polar bear and giant panda (*Ailuropoda melanoleuca*) genomes with 249,740 copies and 237,604 copies, respectively. SINEC1_Ame has a consensus length of 201 bp and was initially described from the giant panda genome (Li et al. 2010). SINEC elements are thought to be LINE1 propagated, and a screen for potentially active full-length LINE1s revealed 535 copies with two intact open reading frames (ORF) in the polar bear genome. The U6 snRNA that has been strongly associated with LINE1 activity in mammalian genomes (Doucet et al. 2015), was found in 67 copies in the polar bear genome sequence. Repeat landscapes of both polar bear and giant panda genomes indicate the presence of low divergent and thus recently active SINEs (**Supplementary Fig. 3**).

### The TeddyPi pipeline

The TeddyPi pipeline is a modular framework to process TE and SV calls and to prepare datasets for phylogenetic inference. It is written in Python and utilizes established code libraries for biological computing. Parameters and the filter pipeline are configured with comprehensively structured configuration files and allow to create tailor-made filter pipelines for a variety of variant callers. The first module (teddypi.py) processes each sample genome individually and filters the output of the selected variant callers. Several filters and merge-functions are included in this module, and a flexible codebase allows implementation of new functions with little programming knowledge. In the same module, large deletions are transformed to reference-insertion calls on the basis of annotated TEs in the reference genome. It is also possible to make intersections or create non-redundant datasets of the input data in this step. In the second module (tpi_ortho.py), TE insertion data is combined across a set of samples (typically different taxa) to generate presence/absence matrices for Ref+ and Ref− separately. Finally, in tpi_unite.py both matrices are merged to a comprehensive presence/absence matrix that can be exported in tabular-text and NEXUS format. A flowchart of the pipeline is shown in Supplementary Figure 4. TeddyPi is open source and can be accessed on https://github.com/mobilegenome/teddypi. Easy configuration and a modular architecture makes it convenient to adapt TeddyPi to process data from a broad range of TE/SV callers or other integration pipelines such as SVMerge or McClintock (Wong et al. 2010; Nelson et al. 2016). TeddyPi can be applied to any group of organisms where accurate TE/SV calling is feasible.

**Figure 1.**
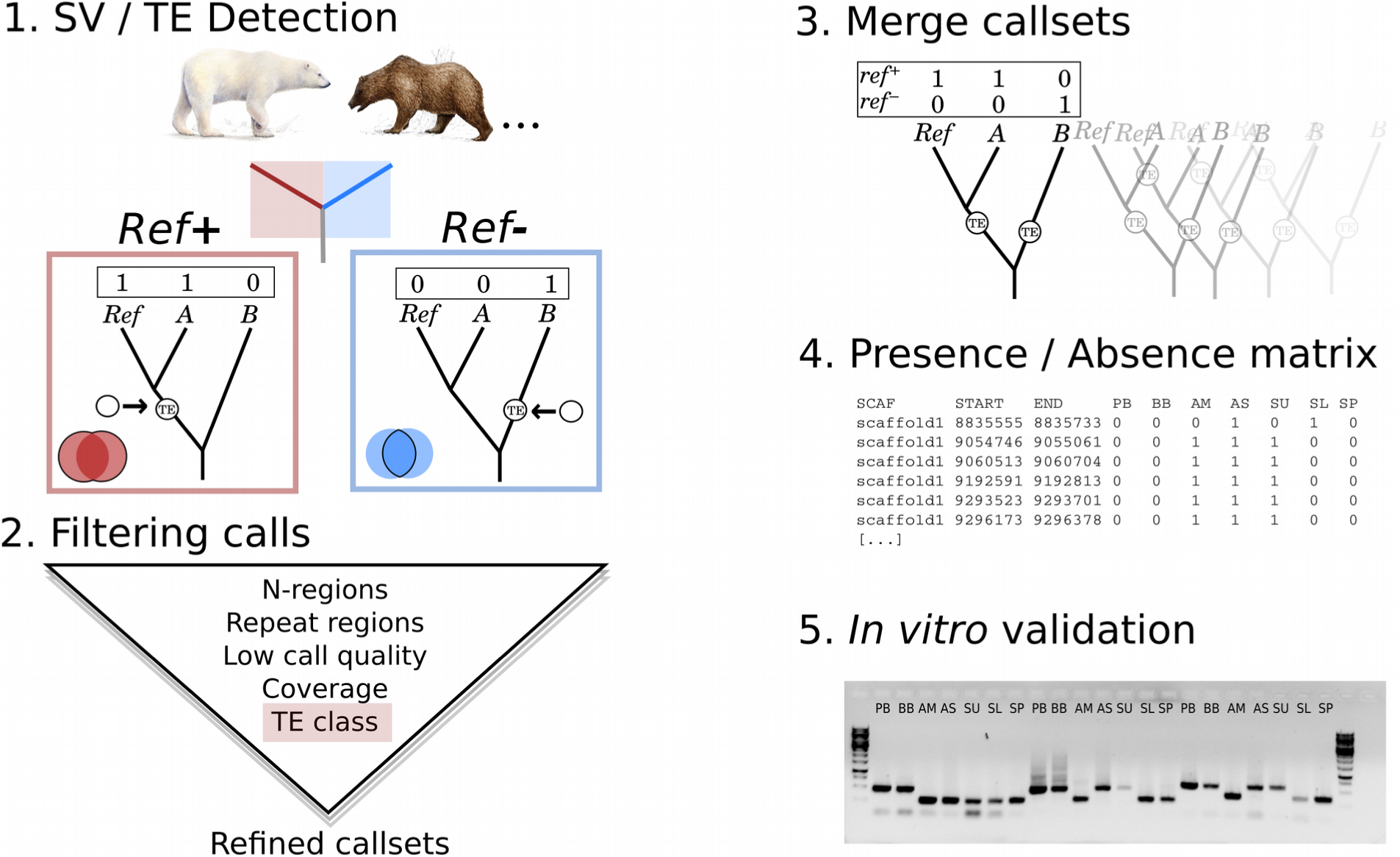
Schematic illustration of the TeddyPi pipeline. 1. Transposable Element (TE) and Structural Variation (SV) callers detect reference (Ref+, red) and non-reference (Ref-, blue) TE insertions from reads mapped to a reference genome. The boxed trees show a schematic phylogeny with the reference genome (Ref) and two other taxa (A and B). The TE insertion is shown by an arrow and indicates Ref+ and Ref− detection depending on which branch the TE inserted. 2. TE calls are filtered based on the polar bear genome annotation, call quality, and sequencing coverage across the genome. Different TE classes are collected separately. 3. Sets of TE calls (call sets) for each individual genome are merged to create a comprehensive presence/absence matrix (4) that is used for phylogenetic inference and (5) to select loci for *in vitro* validation.

### Detecting Ref-insertions

In all analyzed samples, the programs RetroSeq (Keane et al. 2013) and Mobster (Thung et al. 2014) found 696,041 and 491,193 Ref− TE insertions, respectively (**Supplementary Table 4, Supplementary Table 5**). Despite the difference in number of raw calls, the number of SINEs and LINEs selected from the unfiltered datasets of RetroSeq and Mobster are very similar (∼300,000 SINEs, ∼135,000 LINEs). Still, they differed in susceptibility to the subsequent filtering pipeline, indicating differences in overall call-quality (**Supplementary Table 4, Supplementary Table 5, Supplementary Table 6, Supplementary Table 7**). Thus, after filtering 50% more SINEs were obtained from Mobster than from RetroSeq; for LINEs 25% more calls from RetroSeq were retained (**Supplementary Table 8**). The final dataset consisted of 84,462 SINEs and 7,734 LINEs with merged data from RetroSeq and Mobster (**Supplementary Table 8**).

### Detecting Ref+ insertions

A different approach was necessary to identify Ref+ TE insertions, i.e. those shared with the polar bear due to its nested position among the ursine bears (Fig. 1, **Supplementary Fig. 1**). The two SV callers Pindel (Ye et al. 2009) and BreakDancer (Chen et al. 2009) identified in total 10,527,959 deletions in the nine bear genomes of these 96.4% were shorter than 100 bp and excluded. Length distributions of the deletion callsets showed distinct peaks of 200 bp and 6 kb, corresponding to full-length copies of SINEs and LINE1s, respectively (**Supplementary Fig. 5, Supplementary Fig. 6**). After filtering, we retained 12,865 (Pindel) and 296,013 (BreakDancer) high-quality deletion calls that were between 100 bp and 10 kilo basepairs (kb) long (**Supplementary Table 9, Supplementary Table 10**).

The majority (95%) of detected Pindel deletions were also identified by BreakDancer, suggesting a higher reliability at the expense of lower sensitivity in the program Pindel. The filtered data of both programs were merged into a non-redundant set of 295,434 deletion calls (**Supplementary Table 11**). Of these, 270,689 (92%) matched TE annotations in the polar bear genome, and hence were considered as Ref+ TE insertions. We detected 210,999 deletions that intersected SINE insertions in the polar bear genome. From 30,609 deletions matching LINE1 insertions, only a minor fraction (2.5%) was longer than 5 kb, the remaining copies were likely 5’-truncated (**Supplementary Table 11**).

Phylogenetic networks generated from Ref+ and Ref− datasets respectively show that one type of detected insertion can only resolve one side of the tree (**Supplementary Fig. 7, Supplementary Fig. 8**).

### TE insertion rates in ursine bears

For both Ref+ and Ref-insertions, TeddyPi discovered on average 10,000 and 20,000 TE insertions per genome (Fig. 2a). The few TE insertions discovered in the two re-sequenced polar bears reflect the species’ low genetic diversity and are expected because the reference genome is a conspecific. Compared to LINE1 insertions, novel SINE insertions were approximately 6-fold more frequent and 50% more Ref+ than Ref− insertions were identified in the bear genomes (Fig. 2a). The highest number of TE insertions was found in the spectacled bear and lowest number of TE insertions was identified in the two additional polar bear genomes (Fig. 2a). For the other species, the numbers of identified TE insertion were homogeneous. As expected from their higher abundance, the genomic distance between SINE insertions was shorter than for LINEs (median distance: 10,010 bp and 73,240 bp, respectively (Fig. 2b). The upper bound of the LINE1 distances of more than 1 Mb indicates the presence of large genomic regions that are devoid of ursine-specific LINE1 insertions.

**Figure 2.**
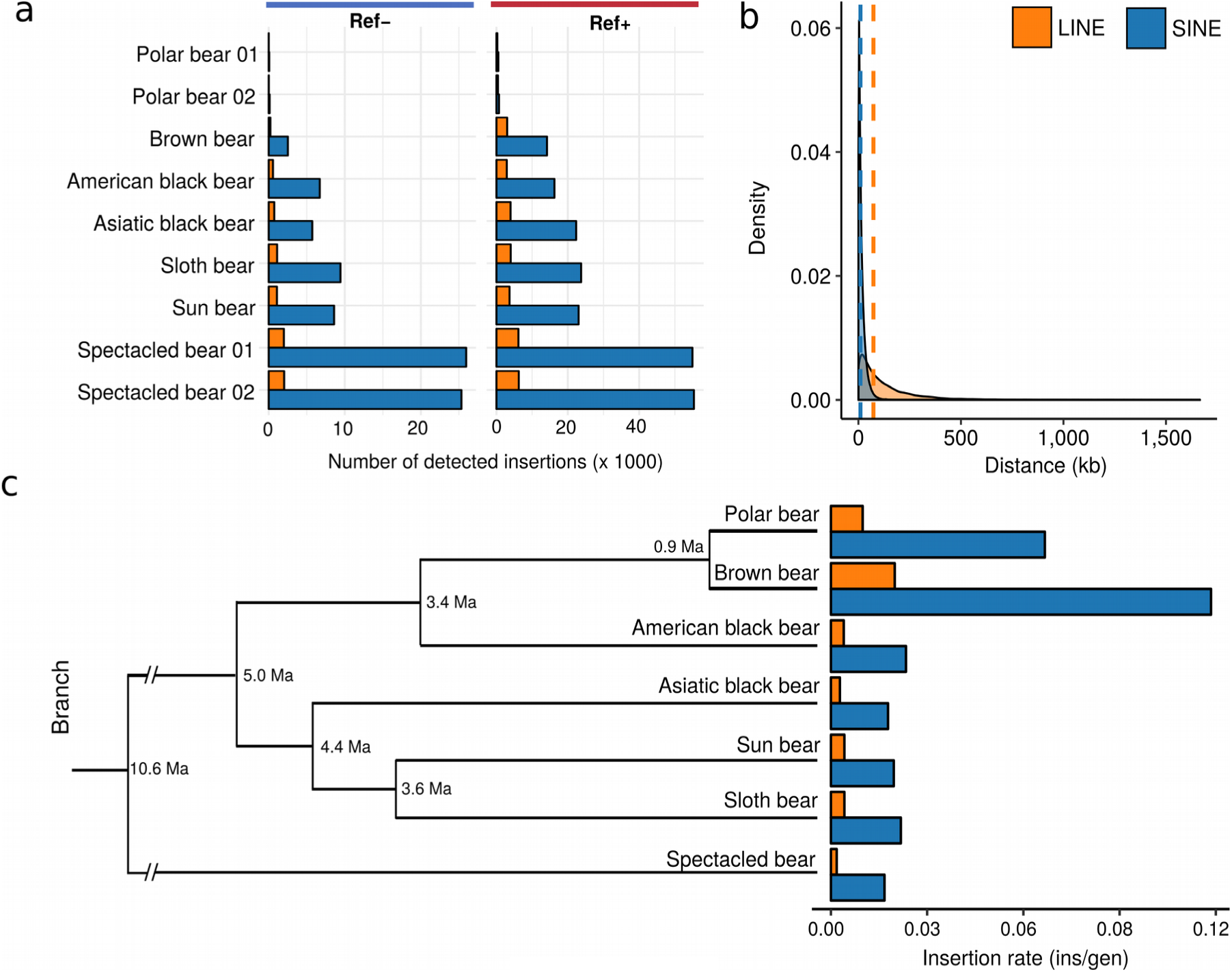
Detection results for TE insertions calls and inferred TE insertion rates. a) Counts of Ref− (left) and Ref+ (right) TE calls per analyzed sample shown for long interspersed element (LINE) insertions (orange) and short interspersed element (SINE) insertions (blue). b) Distance distribution of all detected TE insertion among all bears. Vertical dashed lines indicate median distances. c) TE insertion rates as insertions per generation (ins/gen) for all ursine species were estimated for the terminal branches in a chronogram scaled to divergence times from Kumar et al. (2016).

The rate of TE mobilization is known to differ between lineages (Hormozdiari et al. 2013). Among bears, LINE1-mediated retrotransposition of LINEs and SINEs is ubiquitous, but insertion rates (i.e. number of TE insertion per generation) were substantially higher in brown and polar bear (Fig. 2c). With 0.12 SINE insertions per generation, the insertion rate in the brown bear was the highest. TE insertions into coding or regulatory regions disrupt reading frames or inhibit transcription, however beneficial and potentially adaptive TE insertions are known (Cordaux and Batzer 2009; Casacuberta and Gonzalez 2013; Hof et al. 2016). In bears, 97% of TE insertions integrated into non-coding regions and only few are located in exons or potentially regulatory regions (**Supplementary Fig. 9**).

### In vitro validation of the TE prediction accuracy

Predicting TE insertions from high-throughput sequencing data is challenging and prone to artifacts. We extracted 151 loci to perform validation assays using PCR and Sanger sequencing to assess the accuracy of the *in silico* predictions (**Supplementary Data 1**). All Sanger-sequenced loci where the size of the PCR amplicon suggested a TE insertion were validated as a SINEC1_Ame insertion. Furthermore, the target site duplication (TSD) and breakpoints were identical among bears indicating a single, unique integration event (**Supplementary Fig. 10, Supplementary Note 1**). When compared to the *in silico* predictions, Ref− TE calls were confirmed to 90%, with a false-positive rate (FPR) of 4% and a false-negative rate (FNR) of 6% (Table 1). The results indicate that the Ref− callers are more likely to miss a true TE insertion, than to return an artifact. Loci were randomly selected for PCR validation from the whole dataset or predefined presence/absence patterns for phylogenetic hypotheses (**Supplementary Table 2**). Irrespectively, of whether the hypothesis matched the species tree or is in conflict with it, 93% of the predictions were experimentally confirmed to be accurate (Table 1, Supplementary Data 1).

In all 40 verified Ref+ TE insertion loci an insertion was present in the polar bear, proving the reliability of our approach to select for Ref+ TE insertions. Prediction accuracy for Ref+ insertions in other species was 74%, and was mainly attributed to a higher FPR than in Ref− insertions. A false positive Ref+ TE insertion call, means that deletions were not recovered by SV callers, therefore Ref+ FPR should be considered as FNR. For 111 loci, the PCR amplification yielded an unambiguous phylogenetic informative signal, i.e. amplicon size differences with amplification success in all species. For 40 additional loci, one or more individual did not yield a PCR amplicon, and the locus was recorded as inconclusive for reasons of stringency. For all in vitro validated loci, we identified 17 loci with heterozygous SINE insertions (**Supplementary Table 12**). In the brown bear, 17% of the amplified insertions were heterozygous. For the American black, Asiatic black, sun, sloth and polar bear TE heterozygosity was 6% or less.

**Table 1:** Summary of *in vitro* TE validation experiments for Ref− and Ref+ insertion loci. Results are shown for loci that were phylogenetically informative and all loci, i.e. those lacking amplicons in more than one sample (All). The number of tested loci (N) and frequency of amplicon size differences that matched the computational prediction (true positives, TP), and false positively (FP) or false negatively (FN) predicted insertions are shown. For Ref− loci, random loci (Random), and loci predicted to support a specific phylogenetic hypothesis (Hypothesis-driven) were selected. For Ref+ markers, all loci were randomly selected.

| Type | Set | Informative loci |  |  |  | All loci |  |  |  |
| --- | --- | --- | --- | --- | --- | --- | --- | --- | --- |
|  |  | N | TP | FP | FN | N | TP | FP | FN |
| Ref- | All Ref- | 80 | 0.90 | 0.04 | 0.06 | 111 | 0.87 | 0.07 | 0.06 |
|  | Hypothesis-driven | 48 | 0.93 | 0.03 | 0.04 | 71 | 0.88 | 0.05 | 0.07 |
|  | Random | 32 | 0.82 | 0.05 | 0.06 | 40 | 0.80 | 0.13 | 0.07 |
| Ref+ | All Ref+ | 31 | 0.74 | 0.23 | 0.04 | 40 | 0.70 | 0.26 | 0.03 |
|  | Pindel + BreakDancer | 17 | 0.76 | 0.23 | 0.02 | 20 | 0.71 | 0.28 | 0.01 |
|  | Pindel | 8 | 0.79 | 0.14 | 0.07 | 10 | 0.70 | 0.24 | 0.06 |
|  | BreakDancer | 6 | 0.67 | 0.31 | 0.02 | 10 | 0.71 | 0.27 | 0.14 |

Interestingly, two SINE insertions (No. 40 and 122) were present in all ursine species except the polar bear. The flanks around the empty insertion site in the polar bear lack deletions and only the pre-integration site is present compared to the other ursine bears. Other validated species-tree incongruent TE insertions (**Supplementary Fig. 11**) support alternative tree topologies reflecting the species tree supported by mitochondrial data or previously identified gene-flow signals from individual gene trees (Yu et al. 2007; Kutschera et al. 2014; Kumar et al. 2016). For example, seven validated TE insertions are synapomorphic for American and Asiatic black bear and nine insertions are shared by Asiatic black bear and sloth bear.

### Reconstructing the phylogeny of bears

The Ref+ and Ref− TE insertions were merged into a common dataset that included 150,513 SINE and LINE1 insertions. From these, 71,444 (47.5%) of the TEs were phylogenetically informative and 46.7 % (70,356) were species-specific. We found 8,713 TE insertions being shared by all seven bear species. However these numbers differ when applying maximum parsimony that accounts for missing data (Fig. 3).

We identified seven times more insertion of SINEs than LINEs (132,093 and 18,420, respectively). The phylogenetic analysis focused on SINE insertions because these are shorter than the mean insert-size of the sequencing libraries and thus robustly recovered by TE and SV calling. Dollo parsimony analysis of 132,093 SINE insertions resulted in a phylogenetic tree with 100% bootstrap support for all nodes, except the node separating the two polar bear individuals (Fig. 3). The tree clearly groups spectacled bears that belong to the family Tremarctinae, outside the ursine bears. Within Ursinae, the tree has two clades that consist of the polar, brown and American black bear and the Asiatic black, sun and sloth bear, respectively. Sun and sloth bear form a sister group to the Asiatic black bear. Despite, having 100% bootstrap support and branches that are generally supported by more than thousands of independent SINE insertions, a rescaled consistency index of 0.567 indicated phylogenetic incongruence among the data.

**Figure 3.**
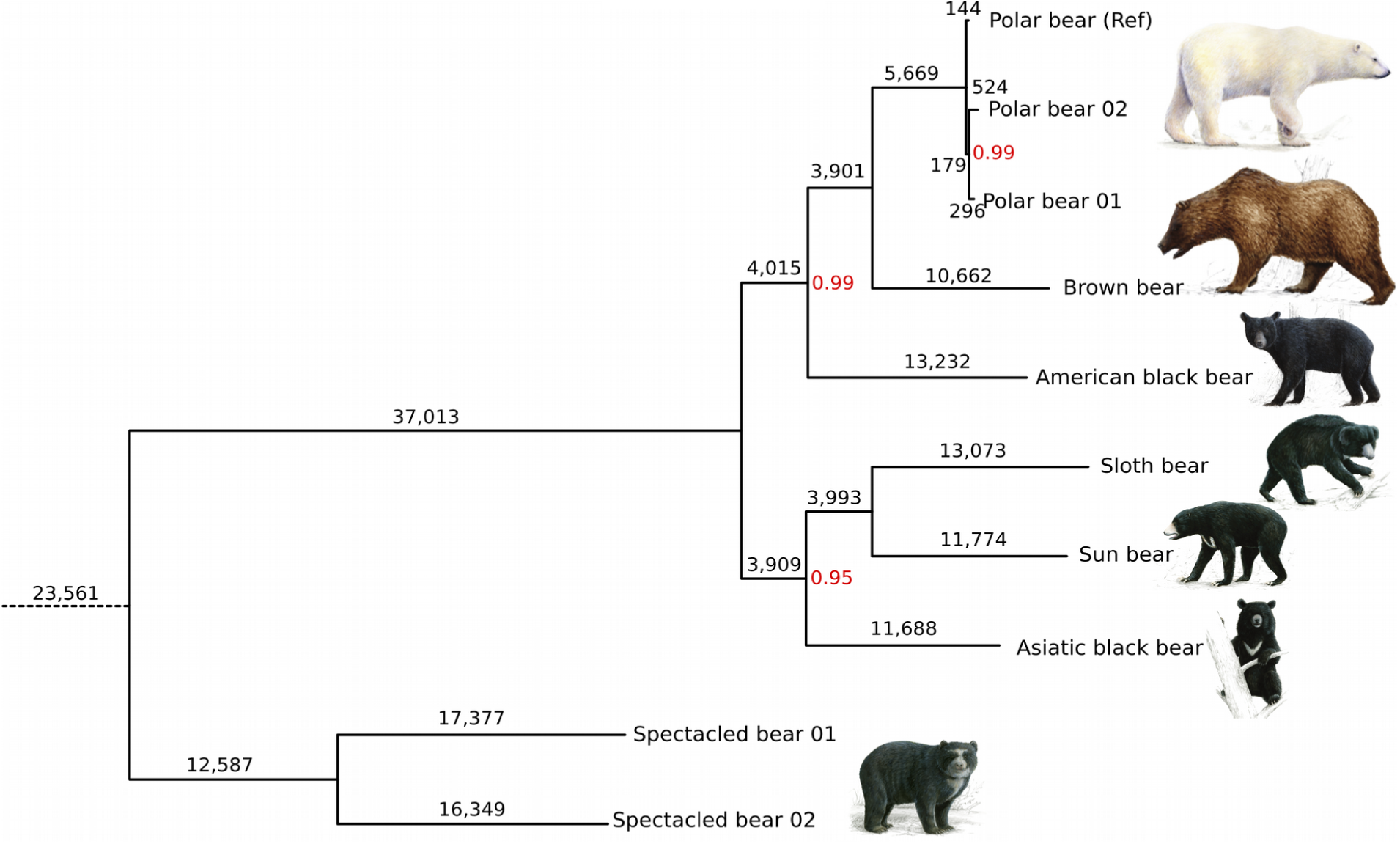
Dollo-parsimony tree of bears reconstructed from 132,039 SINE insertions. Branch lengths indicate the number of SINE insertions on that branch. Most nodes received bootstrap support of 100% (not indicated). Bootstrap support below 100% is shown in red. The rescaled consistency index is 0.567, indicating conflict in the dataset.

To explore phylogenetic conflict, a network analysis of the same data revealed a tree-like network, that clearly separated the Asiatic black, sloth and sun bear from the other three ursine bears by a long edge representing 3,305 SINE insertions (Fig. 4). Still, strong conflict among the Asiatic black, sun and sloth bear was indicated by an intertwined web between them, that also included common splits with polar or brown bear. Polar and brown bear were grouped by an edge that represents 3,597 SINE insertions, but polar bears also shared 2,240 insertions with the American black bear.

Phylogenetic conflict can be caused by hybridization or ancient polymorphisms that lead to allele sharing between non-sister group lineages and has been demonstrated for different ursine bears (Kutschera et al. 2014; Kumar et al. 2016). We analyzed the phylogenetic conflict among Asiatic black, sun and sloth bear using shared SINE insertions obtained from the presence/absence matrix. The Asiatic black bear shares 278 SINE insertions with the sun bear and 265 SINE insertions with sloth bear. The monophyly of sun and sloth bear is supported by 168 SINE insertions. For these three taxa, statistical analyses using the KKSC-test (Kuritzin et al. 2016) support the species-tree topology at high significance (bifurcation test, p=2.325e-10) and reject hybridization between sun bear and the Asiatic black bear (hybridization test, p=0.6060, **Supplementary Table 13**). For American and Asiatic black bear, 129 shared SINE insertions were recovered (Fig. 5b), however the statistical significance of this result could not be assessed with existing methods. The monophyly of polar and brown bear is supported by 3,160 SINE insertions and the species-tree topology of polar, brown and American black bear share is significantly supported (tree test, p=1.04e-159). All three species share 2,178 SINE insertions (**Supplementary Fig. 12**).

**Figure 4.**
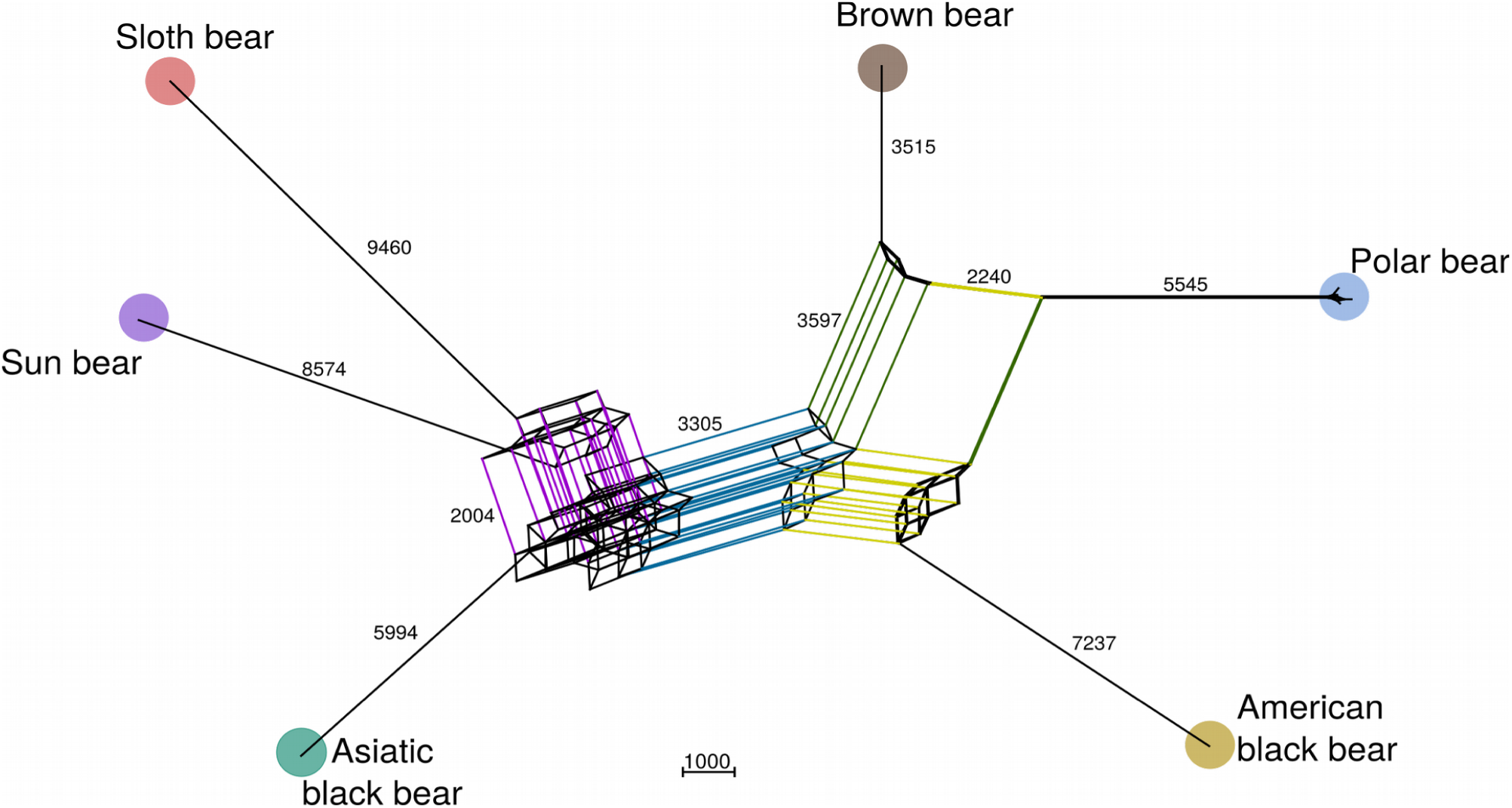
Median network from 132,093 SINE insertions. Parallel edges indicate shared splits between species. Major edges are colored, they separate the two major ursine clades (blue), or group together sun and sloth bear (purple), brown bear and polar bear (green) and American black and polar bear (yellow). Edge lengths indicate the number of shared SINE insertions as calculated by SplitsTree 4. For better readability the spectacled bear is not shown.

**Figure 5.**
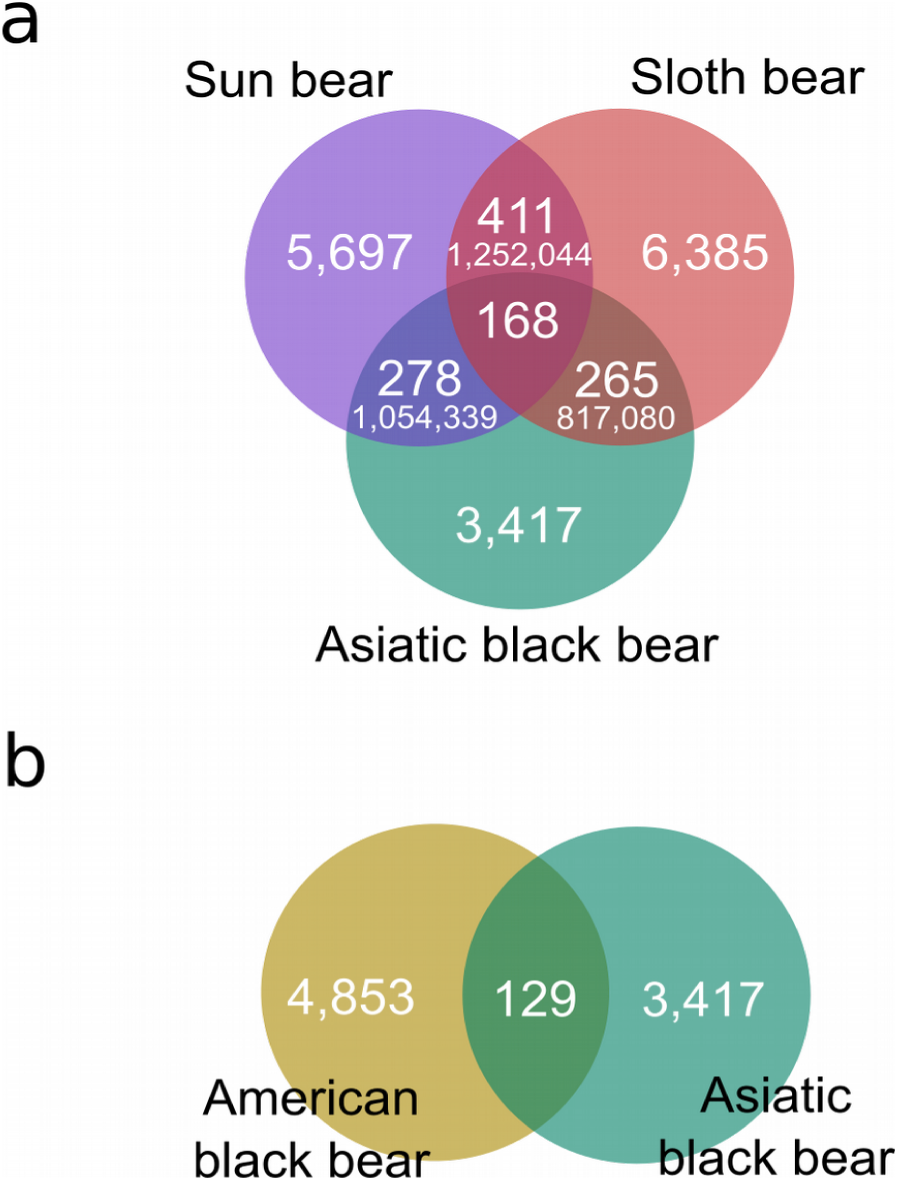
Venn Diagrams depicting phylogenetic conflict among Asiatic black, sun and sloth bear (a) and American black and Asiatic black bear (b). The amount of shared SINE insertions under Dollo-parsimony are shown. The numbers in smaller font (a) give the amount of shared nucleotide substitutions (Kumar et al. 2016)

### Different extent of phylogenetic signal in the flanking regions

Alignments of genomic sequences flanking phylogenetically informative TE insertion sites were analyzed for their phylogenetic signal and if it is congruent with the phylogenetic signal from the corresponding TE insertion. Up to 65% of the individual maximum likelihood (ML) trees calculated from the flanking sequences reconstructed were identical with the presence/absence pattern of the TE insertion (Fig. 6). To investigate the spatial congruence between the TE insertion and its flanks in more detail, we measured the number of substitutions reconstructing the same phylogeny as the TE insertion in 1 kb non-overlapping windows extending up to 10 kb from the insertion site (Fig. 6). TE supporting substitutions were elevated in direct vicinity of the TE insertion site and then tapered off with distance from the insertion site. Also, the frequency of supporting substitutions is highest at TE insertion sites, that are congruent with the ursine species tree and lower for those with conflicting signal. For example, among 215 orthologous TE insertions shared by all Asiatic bears, the average frequency of TE-supporting substitutions increased from 0.01 to 0.04 within the first 5 kb from both sides of the insertion site (Fig. 6). For species-tree incongruent TE insertion loci, elevation of TE-supporting substitutions was less pronounced and the stretch of spatial congruence was shorter. Substitution frequencies for phylogenies that are different to the TE insertion signal were generally not elevated towards the insertion site (**Supplementary Fig. 13**). In cases of minor difference in phylogenetic signal between substitutions and TE, substitution frequencies were raised despite of different signals (**Supplementary Note 2**).

**Figure 6.**
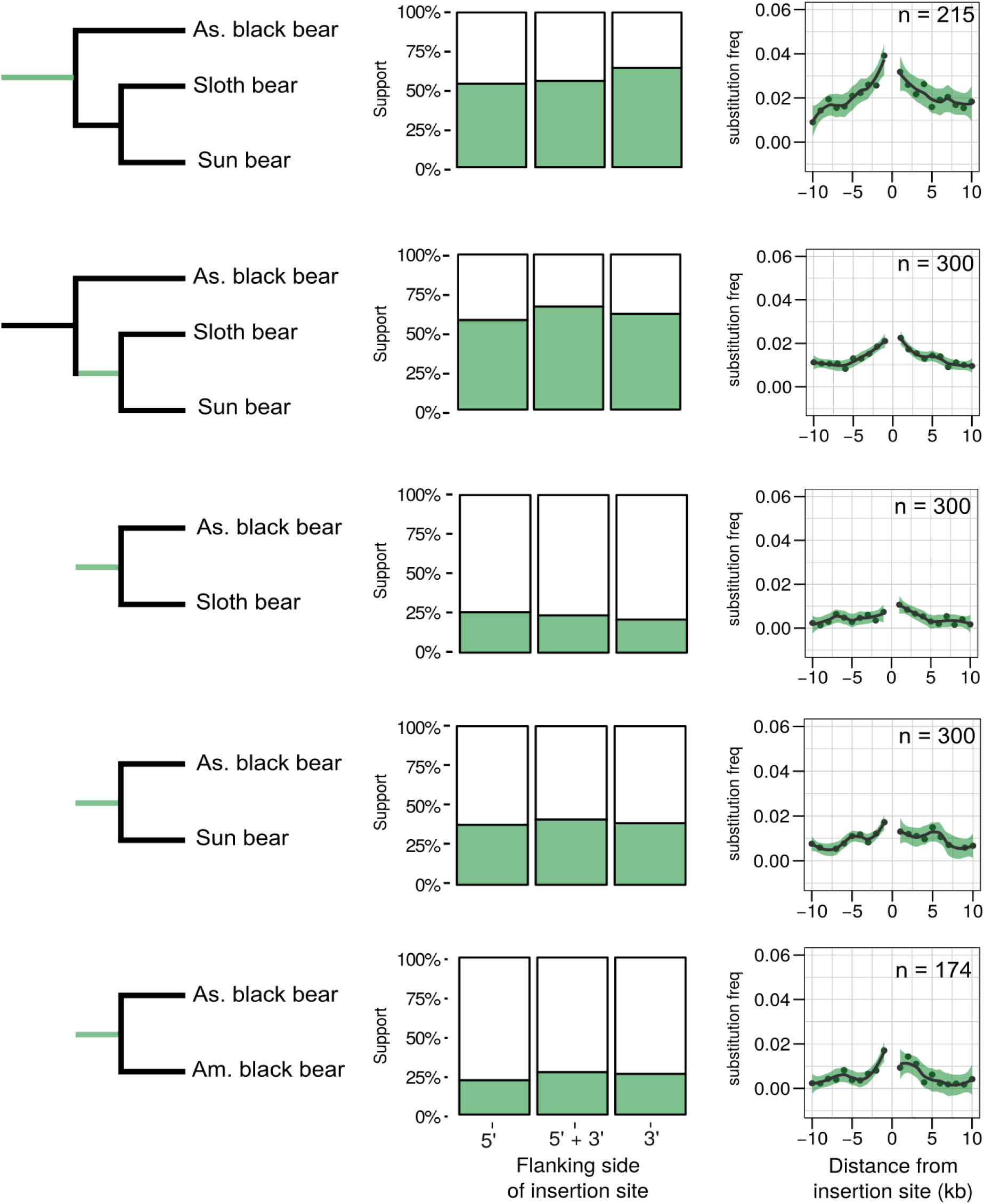
Analysis of flanking sequences of TE insertions present in different groups of taxa. Left panel: Green branches in the phylogenetic tree indicate when the TEs integrated. Middle panel: Bar plots showing the frequency of ML-trees calculated from 10 kb flanking sequence on the 5’, 3’ end or a concatenation of both. Left panel: Frequency of substitutions that support the TE insertion signal in 1 kb windows around the insertion site. Frequencies are normalized by the number of segregating sites.

## Discussion

Analyzing whole genome sequence data for TE insertions allows studying the landscape of genetic variation at unprecedented extent and detail. However, it faces methodological challenges. Here, we developed the TeddyPi pipeline that integrates different available TE callers and applies stringent filtering to overcome limitations of TE calling. It also produces an automated output of presence/absence tables of TE insertions that can be immediately used for phylogenetic analyses. The pipeline follows a ‘quality over quantity’ approach to select only highly reliable TE insertion loci. Recent phylogenomic studies suggest that genomes are often a mosaic of different genealogies caused by evolutionary processes such as introgressive hybridization or ILS (Mallet et al. 2016). To study such complex signals, sufficient character sampling is necessary. This can only be achieved by nucleotide-based genome analyses, or genome-wide and ascertainment bias free discovery of TE insertions (Kuritzin et al. 2016; Dodt et al. 2017). TE insertion data provide an independent and robust molecular marker system to build phylogenies that are not based on sequence analysis (Shedlock et al. 2004).

### SINE insertions recapitulate the evolutionary history of bears

Extensive phylogenetic discordance across loci has previously challenged the resolution of the bear phylogeny (Yu et al. 2007; Kutschera et al. 2014; Kumar et al. 2016). The TeddyPi pipeline extracted more than one 100,000 TE insertions from low-coverage data to build a reliable dataset of phylogenetically informative TE markers to study the evolutionary history of bears. We reconstructed a well-supported phylogenetic species tree despite incongruent phylogenetic signals (Fig. 3, Fig. 4). The three Asian bears form a clade that is consistent with coalescent analyses of genome sequence data (Kumar et al. 2016). However, this contrasts previous studies, that placed the Asiatic black bear as sister group to the polar, brown and American black bear clade or as sister group to the American black bear, respectively (Yu et al. 2007; Krause et al. 2008; Pagès et al. 2008). Despite significant bootstrap support for each node of the parsimony TE tree, the tree had a low consistency index, indicating that many TE insertions conflict with the inferred phylogeny. Phylogenetic networks can depict such conflicting signals better than trees that force the data to a bifurcating model of evolution (Bapteste et al. 2013). The network analyses reveals that phylogenetic conflict among bears occurs mostly in the two main clades of the ursine subfamily (Fig 4). In particular, the Asiatic black, sun and sloth bear that currently inhabit South-East Asia form a complex network. We explored this conflict further and found that the Asiatic black bear share almost identical numbers of orthologous SINE insertions with sun and sloth bear respectively, thereby indicating ILS as origin of the conflict (Fig. 5, **Supplementary Table 13**). Despite reconstructing the same species tree, our detailed analyses contrasts nucleotide-based analyses of millions of sites, that inferred ancestral hybridization as main driver of phylogenetic conflict among these species (Kumar et al. 2016). To what extent hybridization occurred between bears and what caused the conflicting signal of single nucleotide substitutions and TE insertions remains to be further explored.

In previous mtDNA-based analyses the Asiatic and American black bear have been placed as sister species(Yu et al. 2007; Krause et al. 2008). This is not supported by the majority of identified TE insertions. However, 129 SINE insertions are shared by American and Asiatic black bear (Fig. 5). Therefore, the close relationship of the two black bears based on mtDNA is likely a result of an ancient mitochondrial capture event and additional introgression of nuclear DNA carrying these TE insertions (Kutschera et al. 2014). An alternative scenario explaining the discordance between mtDNA and nuDNA phylogenies of American and Asiatic black bear involves nuclear swamping of the American black bear genome by brown bear alleles. This would produce a similar phylogenetic signal and artificially place the American black bear on the lineage leading to brown and polar bear (Kutschera et al. 2014). However, our network analysis and 99 shared SINE insertions by brown bear and American black bear yields very little support for this hypothesis suggesting that ancient hybridization between the two black bear species had a more pronounced effect on their genomes than nuclear swamping by brown bear DNA (Fig. 3, **Supplementary Fig. 12**).

Differences in retrotransposition activity or demographic history can cause varying rates of TE insertion to between lineages (Hormozdiari et al. 2013). The insertion rates were estimated to 0.022 SINE and 0.004 LINE1 insertions per genome per generation, which is half of the rate for humans (0.035 Alus and 0.008 LINE1s) (Fig. 2c, Sudmant et al. 2015). Fixation of neutral or slightly deleterious TE insertions depends on genetic drift, that is stronger in small effective population sizes or on purifying selection, which is stronger in large populations (Charlesworth 2009; Gonzalez and Petrov 2012). Substantially higher insertion rates of TEs and a high heterozygosity rate in brown bear thus can be explained by large population size that brown bears maintained over long timespans (Miller et al. 2012). The high TE insertion rate in polar bear is unexpected given its low genetic diversity (Hailer et al. 2012). Also, insertions of mitochondrial sequences appear to be less frequent in bears than in other species (Lammers et al. 2016), making it necessary to explain the elevated rate of TE insertions. A possible explanation would be retrotranspositional burst caused by hybridization (O’Neill et al. 1998; Dion-Côté et al. 2014). Bears in general can hybridize, and hybrids between polar and brown bears have been observed (Galbreath et al. 2008; Kelly et al. 2010). Additionally, a hybrid origin of polar bear has been proposed (Lan et al. 2016). Thus, consequent genetic introgression potentially lead to a burst of TE insertions in the species into which hybrids backcross and thus may explain the high TE insertion rate in brown and polar bears.

The accompanying sequence-based analyses of the same dataset enabled to examine the correlation of nucleotide substitutions and TEs for conflicting phylogenies (Kumar et al. 2016). Expectedly, TE insertions were several magnitudes less frequent than nucleotide substitutions. Yet, both analyses yielded the same phylogeny but differed in their interpretation of phylogenetic conflict (Fig. 5). This highlights the need for nucleotide-based analyses in addition to genome wide analyses of TE insertions.

### Quality over quantity approach for phylogenetic inference of TEs

Previous phylogenetic TE analyses relied on the availability of reference genomes which were often restricted to one species per order or family. For bears, draft genome assemblies of polar bear and giant panda are available (Li et al. 2010; Liu et al. 2014) and traditional *in vitro* approaches would have identified orthologous loci in both genomes, with one carrying a TE insertion that is experimentally tested using PCR in the other bear species for presence or absence (Shedlock et al. 2004). Although availability of two reference genomes is beneficial, unbiased identification of variable i.e. phylogenetically informative TEs across the complete taxon-sampling, is not possible this way. Adding genomes from the entire ursine subfamily enables discovery of TE insertions that is free from sampling artifacts and allows targeted extraction of phylogenetically informative markers. However, the nested position of the polar bear reference genome inside the species tree, the use of low-coverage genome data and misassembled regions in the reference genome is challenging and required methodological refinements to increase prediction quality of TE insertions. These challenges were rarely discussed in other studies but are central when aiming for a large-scale identification of TE insertions from paired-end mapping data without introducing a sampling bias.

If the reference genome is nested inside the ingroup, as in the case of the polar bear inside Ursinae, a two-sided approach using Ref+ and Ref− insertions is necessary to yield unbiased support for all internodes in the resulting phylogenetic tree or network (**Supplementary Fig. 2**). The polar bear genome sequence has higher contiguity than that of giant panda, has a better assembly of repeats due to longer sequencing reads and it benefits from the low heterozygosity in polar bear. Also, the giant panda is less ideal to be used as reference genome for mapping because of its high evolutionary distance to the other bear species, which diverged from the giant panda some 20 Ma. On the other hand the polar bear is nested within the other bears, which makes variant calling more difficult. To solve this problem and to make TeddyPi more ubiquitously applicable, SV callers were integrated in the pipeline to deduce Ref+ insertions from deletions calls (Nellåker et al. 2012). Only few TE callers are specifically developed to detect Ref+ insertions. To our knowledge, only T-lex and T-Lex2 (Fiston-Lavier et al. 2011; Fiston-Lavier et al. 2015) perform Ref+ insertion detection, but they are not compatible with the TeddyPi pipeline due to different file format requirements. Other programs, such as RetroSeq, Mobster and Jitterbug exclusively detect Ref− TE insertions (Keane et al. 2013; Thung et al. 2014; Hénaff et al. 2015). Depending on the mapping-signature utilized for SV-calling (split-reads, read-pairs, depth of coverage) detection results differed markedly between programs as exemplified by our results from Pindel and Breakdancer (**Supplementary Table 9, Supplementary Table 10**) and from other studies (Ewing 2015). Inconsistencies between different programs will affect the phylogenetic inference, which relies on precise presence/absence patterns of orthologous loci and making it necessary to integrate different SV callers as implemented in TeddyPi. While TE calls from Mobster and RetroSeq were almost concordant, still overlapping calls were used to increase the reliability of the calls. For TE calling, integration of multiple callers is recognized as an appropriate strategy to enhance the consistency of TE predictions (Lin et al. 2015; Nelson et al. 2016), and this functionality is implemented in TeddyPi for both, Ref+ and Ref− insertions. A true positive rate (TPR) of 93 % for TE calls from the TeddyPi pipeline (Table 1) is higher than the estimated sensitivity of RetroSeq for 10X whole genome sequencing data (Keane et al. 2013). The reliability of TeddyPi is equally as estimates from Mobster analyses of high-quality human data. Thus, when possible, the use of a suitable outgroup genome to analyze only Ref− insertions for phylogenetic reconstruction is recommended.

Detecting TE insertions and SVs in resequenced whole genome data often have breakpoint inaccuracies within a margin of up to 50 bp (Ewing 2015). It is therefore not possible to distinguish between near or near-exact deletion or insertions. This can affect detecting ortholog events or analyzing genetic effects by intersection with coding sequences (**Supplementary Fig 7**). However, previous studies have indicated that long near-exact indels occur at a very low level and would therefore contribute only marginally to the observed phylogenetic conflict among bears (van de Lagemaat et al. 2005).

Missing data and unplaced scaffolds are common in most genome assemblies, because of current technological limitations to sequence and assemble repetitive DNA. Thus, in genome sequences, sequence gaps are mostly caused by repetitive regions, such as TEs and satellite DNA. Soon, long read sequencing technologies, such as PacBio or Nanopore, will likely alleviate this problem considerably, but it is unlikely that the technology will be used routinely, because of the higher sequencing cost. The 2.3 Gb polar bear genome sequence was based on short read technology and lacks 400 Mb of genomic information, based on an estimated genome size of 2.7 Gb for extant bears(Vinogradov 1998; Krishan et al. 2005; Liu et al. 2014). Another artifact from repetitive DNA in genome sequences, are unassembled regions in the scaffolds (N-regions). TeddyPi utilized 38 Mb of N-regions in the polar bear genome as proxy for poorly assembled regions, and all TE calls in their vicinity were excluded from the analyses. The removal of N-regions greatly increased the success rates in experimental validation and show that this is a necessary step in TE calling, that previously have not been implemented in TE calling studies. Another indicator of assembly quality and thus ability to confidently predict TEs is the mappability of short-reads to the reference genome. Mappability can be assessed by deviations of local coverage depth from the mean coverage. To account for poorly mapped regions, TE calls in regions of exceptionally low and high-coverage were coded as missing data. Another challenge to TE and SV calling comes from the random integration of TEs in the genome. Occasionally, young TEs can randomly integrate in older TE sequences. If both TEs are of the same type, sequence reads will be ambiguously mapped to either the young or old TE. This increases the risk for false positive calls during TE calling. Therefore, TE calls located within annotated TEs of same type were removed in the TeddyPi pipeline to increase the reliability of our phylogenetic markers.

Unlike for the human genome, a generally accepted standard or database of TE insertions does not exists for non-model organisms to compare our results to. Thus, detection sensitivity can only be estimated by experimental approaches. The validation experiments show that compared to standard TE callers, the rigorous approach of the TeddyPi pipline substantially improves TE detection from non-model organism genomes that lack highly curated and well-annotated genome assemblies. For the polar bear genome sequence, every experimentally verified loci were confirmed for the presence of SINEC1_Ame, corroborating the assembly and RepeatMasker annotation for these loci. The presence of TSDs in all analyzed loci further strengthens the TeddyPi approach in identifying true, orthologous TE insertion events.

### TE insertions, flanking sequences and recombination blocks in ursine bears

TE insertions share an evolutionary history with nucleotide substitutionsin their immediate genomic vicinity (Daly et al. 2001). If the TE insertion is neutral, the extent of linkage, i.e. the size of a recombination block that carries the TE depend on the recombination rate and the demographic history of the genomic region (Ellegren and Galtier 2016). In great apes, phylogenetic congruence between the TE insertion and its flanking sequence was used to prove hemiplasy of the TE insertion (Hormozdiari et al. 2013), however nucleotide-homoplasy and uncertainties in tree-reconstruction of the specific regions can mislead such an analysis, especially for longer timescales (Suh et al. 2015). Ursine bears radiated around 5 Ma, which left little time for flanking sequences to be saturated, allowing for nucleotide level comparisons. In bears, TE insertions and their flanking sequences share the same phylogenetic signal, but the extent of spatial congruence (i.e. linkage) is limited to a few kb and differs depending on the phylogenetic signal of the TE (Fig. 6, **Supplementary Fig. 12**). The size of the recombination block, as evident from the extent of spatial congruence (Fig. 6), allows estimating the relative time since the TE insertions. A lesser extent of spatial congruence around the species-tree incongruent TE insertions can be explained by an earlier TE integration and subsequent breakdown of the recombination blocks. TE insertions shared exclusively by American and Asiatic black bear have a narrow extent of spatial congruent substitutions, and thus are older than species-tree congruent TE insertions. If a locus originates from more recent introgression a wider extent of spatial congruence carrying the same phylogenetic signal is expected. The flanks of the orthologous TE insertions in the Asiatic bears share the same phylogenetic signal, and therefore show no homoplasy and suggest that ILS has contributed to the phylogenetic incongruence among these loci. For the Asiatic bears, we propose that ILS is the primary driver of phylogenetic incongruence causing high amounts of pairwise similarities (Fig. 5a, Kutschera et al. 2014) and additionally, hybridization between Asiatic black and sun bear led to an excess of shared alleles between these species (Fig. 5a, Kumar et al. 2016). Under the assumption that the current species tree of bears (Fig. 3) reflects the speciation history, introgressive hybridization involving the American black bear must have occurred. However, in agreement with coalescent-based analyses (Kutschera et al. 2014), analyses of TE insertion patterns and their flanking regions (Fig. 5, Fig. 6) indicate that bear lineages are not yet sorted, thereby confounding introgression analyses. Although sequence analyses of the TE flanking regions were restricted to one taxonomic group, it is evident that analyses of deeper divergences in any taxa will lead to shorter recombination blocks and thus fewer phylogenetic signatures. Thus, screening for flanking substitutions surrounding old TE insertions is likely to be uninformative due to the limited spatial congruence and nucleotide saturation.

## Conclusion

Twenty years after the successful introduction of TE insertions as phylogenetic markers, it is now possible to not only use a few, but thousands of informative loci across the genome to reconstruct phylogenies of complete taxonomic groups. The TeddyPi pipeline enables easy *in silico* TE detection in low coverage genomes and provides virtually homoplasy-free evolutionary information that can be used to understand speciation events. The unbiased detection of TEs is essential for the reliability of phylogenetic results. The conceptual framework of the integrated and stringent approach in TeddyPi allows now analysis of ancestry-informative TEs as a routine procedure in comparative genomic studies. Deciphering recent and complex speciation processes using TE insertions as well as nucleotide substitutions is subject to further analyses and important for our understanding of phylogenetics and speciation (Mallet et al. 2016).

## Acknowledgements

The authors thank Dije Tjwan Thung, The Radboud University Medical Center, for providing an unpublished version of Mobster, Thomas Keane, Wellcome Trust Sanger Institute, for advice in using RetroSeq, and Markus Pfenninger for helpful discussions. We are thankful to Kathinka Schulze and Clara Heumann-Kieser for performing validation experiments. Jón Baldur Hlíðberg (www.fauna.is) painted the bears in Fig 3.

## Author contributions

FL, MN and AJ conceived and designed the study. FL developed TeddyPi and performed the computational analyses. SG and MN coordinated and performed experimental validation experiments. FL and MN wrote the manuscript with input from all co-authors. All authors read and approved the final manuscript.

## Data availability

The final TE dataset, and primers for validation experiments are included as Supplementary Data. TeddyPi is available at https://github.com/mobilegenome/teddypi.

